# Container-less Acoustic Levitation Expands Plasma Extracellular Vesicle Proteome Coverage by Mitigating Size-Dependent Peptide Loss

**DOI:** 10.64898/2026.07.25.740687

**Authors:** Edward Huang, Chang Liu, Kent Hoskins, Yu Gao

## Abstract

Proteomic profiling of plasma-derived extracellular vesicles (EVs) is limited in part by adsorptive loss of peptides and proteins to container walls during sample preparation. Here we apply an automated, environment-controlled acoustic levitation platform (Levcell) to the tryptic digestion of small EVs (sEVs) isolated from pooled breast cancer patient plasma, and compare it directly with digestion in low-bind microcentrifuge tubes. Across three parallel technical replicates per method, container-less digestion identified 309 ± 22 protein groups versus 261 ± 6 for tubes (+18.4%; Welch t-test p = 0.053), with equivalent or better quantitative reproducibility (median CV 12.0% vs 14.4%). The gain was strongly asymmetric: 66 protein groups were recovered only under levitation while 10 were recovered only in tubes (exact McNemar p = 3 × 10^-11^). Peptides recovered exclusively by levitation were longer and heavier than those exclusive to tubes (median 14 vs 12 residues, 1611 vs 1358 Da; p *<* 2 × 10^-6^; Cliff’s δ ≈ 0.19–0.20), whereas the total peptide pools were indistinguishable and mean missed-cleavage rates were equivalent (0.276 vs 0.263, p = 0.41), excluding differential digestion efficiency as an explanation. No systematic difference in hydropathy, isoelectric point or hydrophobic residue frequency was detected. The levitation-rescued sub-proteome was enriched for ribosomal, proteasomal, chaperonin and RNA-binding complexes — canonical sEV luminal cargo (MYC targets 16/19, odds ratio 31.7, q = 5.6 × 10^-8^) — and covered 33 of the 100 ExoCarta reference markers versus 22 for tubes, gaining 14 markers while losing three (a single ezrin/moesin/radixin protein group; McNemar p = 9.8 × 10^-4^). We also report two findings that temper the approach: levitated samples carried an approximately 3.7-fold higher keratin burden, consistent with airborne contamination in an open chamber, and no individual marker showed a significant abundance difference after correction for multiple testing. Container-less processing therefore offers a reproducible gain in sEV proteome coverage attributable to reduced size-dependent peptide loss, provided that contamination control is addressed.

## Introduction

Liquid biopsy has become an established route to noninvasive tumour sampling, longitudinal monitoring and treatment selection^1^. Among blood-borne analytes, circulating tumour-derived proteins report directly on tumour phenotype and dynamics^2^, and mass spectrometry (MS)-based proteomics offers hypothesis-free access to that landscape, with applications across screening, diagnosis, prognosis and therapy monitoring^3-8^. Extracellular vesicles (EVs) — in particular the small EV (sEV) population of 30–150 nm — are attractive inputs for such profiling, because the lipid bilayer protects luminal cargo in circulation and because sEVs compartmentalise cell-derived proteins within the plasma matrix^9^. Following MISEV2023 guidance we use the operational term “small EV” throughout rather than “exosome”, since our isolation does not establish endosomal origin.

The analytical demands are severe. The plasma proteome spans more than ten orders of magnitude in concentration, and the proteins of clinical interest sit far below the albumin- and immunoglobulin-dominated bulk^10,11^. Against this background, a technical loss mechanism that would be negligible for abundant analytes becomes decisive: peptides and proteins adsorb irreversibly to plastic and glass surfaces during preparation, reducing recovery and biasing sequence coverage^12-15^. For low-input workflows this cumulative depletion can determine whether an analyte is detected at all.

Acoustic levitation — trapping a liquid droplet in a standing ultrasonic field — removes the liquid–solid interface during processing and has been coupled to MS proteomics in several proof-of-concept studies using protein standards, peptide digests and low-cell lysates^16-20^. Those studies used comparatively simple matrices; whether the approach retains an advantage in a clinical biofluid, where the dynamic range and the co-isolated background are far more demanding, has not been tested.

Here we address that question directly. Using the automated, environment-controlled Levcell platform^17^ developed in our laboratory, we digested sEVs isolated from pooled breast cancer patient plasma in levitated droplets and, in parallel, in low-bind microcentrifuge tubes, and compared the resulting proteomes by DIAMS. We ask three questions: whether container-less digestion increases proteome coverage in a clinical matrix; whether any gain has the size-dependent signature predicted by an adsorptive loss mechanism; and what the practical costs of open-chamber processing are.

## Results

### Container-less digestion increases proteome coverage with equivalent reproducibility

sEVs isolated from pooled breast cancer plasma were digested in parallel triplicate, either as 5 µL acoustically levitated droplets or in low-bind microcentrifuge tubes, and analysed by DIA-MS on identical gradients. Levitated samples yielded 309.3 ± 21.6 protein groups compared with 261.3 ± 5.9 for tubes, an increase of 18.4% (Welch t-test p = 0.053; Figure 1A). At the precursor level the difference was smaller and not significant (4009 ± 124 vs 3662 ± 266, +9.5%, p = 0.138). Quantitative precision was at least preserved: median protein-level coefficient of variation was 12.0% for levitated droplets and 14.4% for tubes, both within the range generally accepted for clinical assays (Figure 1B).

**Figure 1.**
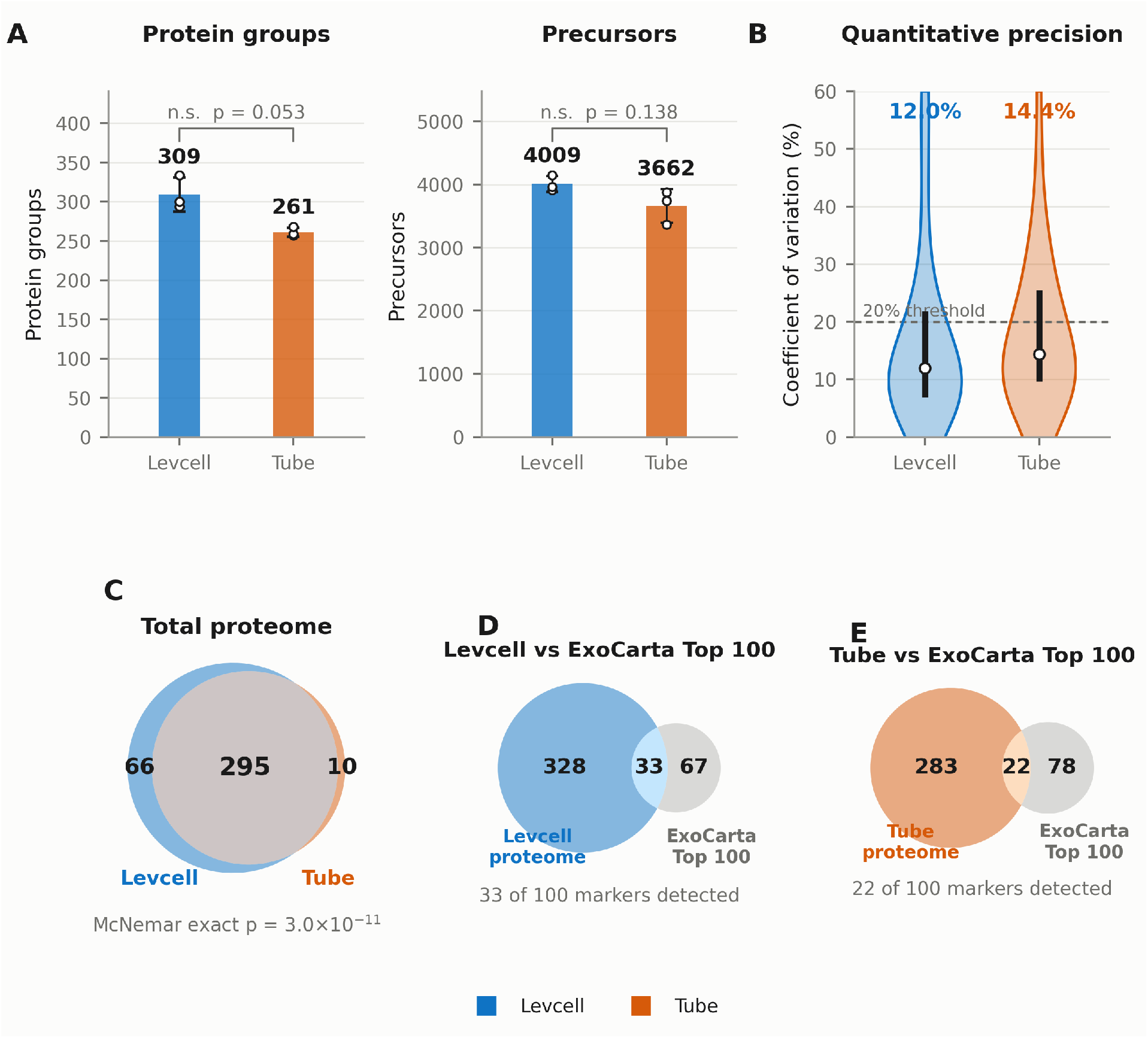
Global proteome coverage and reproducibility. (A) Protein groups and precursors identified per replicate; bars show the mean, error bars the standard deviation, and open circles the three individual replicates. Welch’s t-test; both comparisons are non-significant. (B) Distribution of protein-level coefficient of variation across replicates quantified in all three runs of a workflow; the kernel is clipped at zero, the thick bar spans the interquartile range and the open circle marks the median. The dashed line marks the 20% threshold commonly accepted for clinical assays. (C) Overlap of protein groups detected in at least one replicate of each workflow; the p-value is an exact McNemar test on the discordant counts. (D, E) Overlap of the (D) levitated and (E) tube proteomes with the ExoCarta Top-100 reference marker set. Marker membership is evaluated across all genes listed in each protein group.

Because the two workflows were run on the same input material, the appropriate test of the coverage claim is a paired one on protein presence. Of 371 protein groups detected in total, 295 were common to both methods, 66 were detected only under levitation, and 10 only in tubes (Figure 1C). This 6.6:1 asymmetry is highly significant by exact McNemar test (p = 3 × 10^-11^) and is robust to the presence/absence threshold (requiring detection in ≥ 2 of 3 replicates gives 56 vs 9, p = 2 × 10^-9^). The gain therefore reflects a consistent, directional recovery of additional proteins rather than run-to-run variability.

### The rescued peptide pool is longer and heavier, and is not explained by digestion efficiency

If the additional identifications arise from reduced adsorptive loss, and if adsorption is size-dependent, the rescued analytes should be systematically larger. We compared peptide length and mass across the method-exclusive precursor pools. Precursors detected only under levitation (n = 604) had a median length of 14.0 residues and mass of 1611.3 Da, versus 12.0 residues and 1357.7 Da for those detected only in tubes (n = 322); both differences were highly significant (Mann–Whitney p = 6.6 × 10^-7^ and 1.4 × 10^-6^) with small effect sizes (Cliff’s δ = +0.198 and +0.192) (Figure 2C–F).

**Figure 2.**
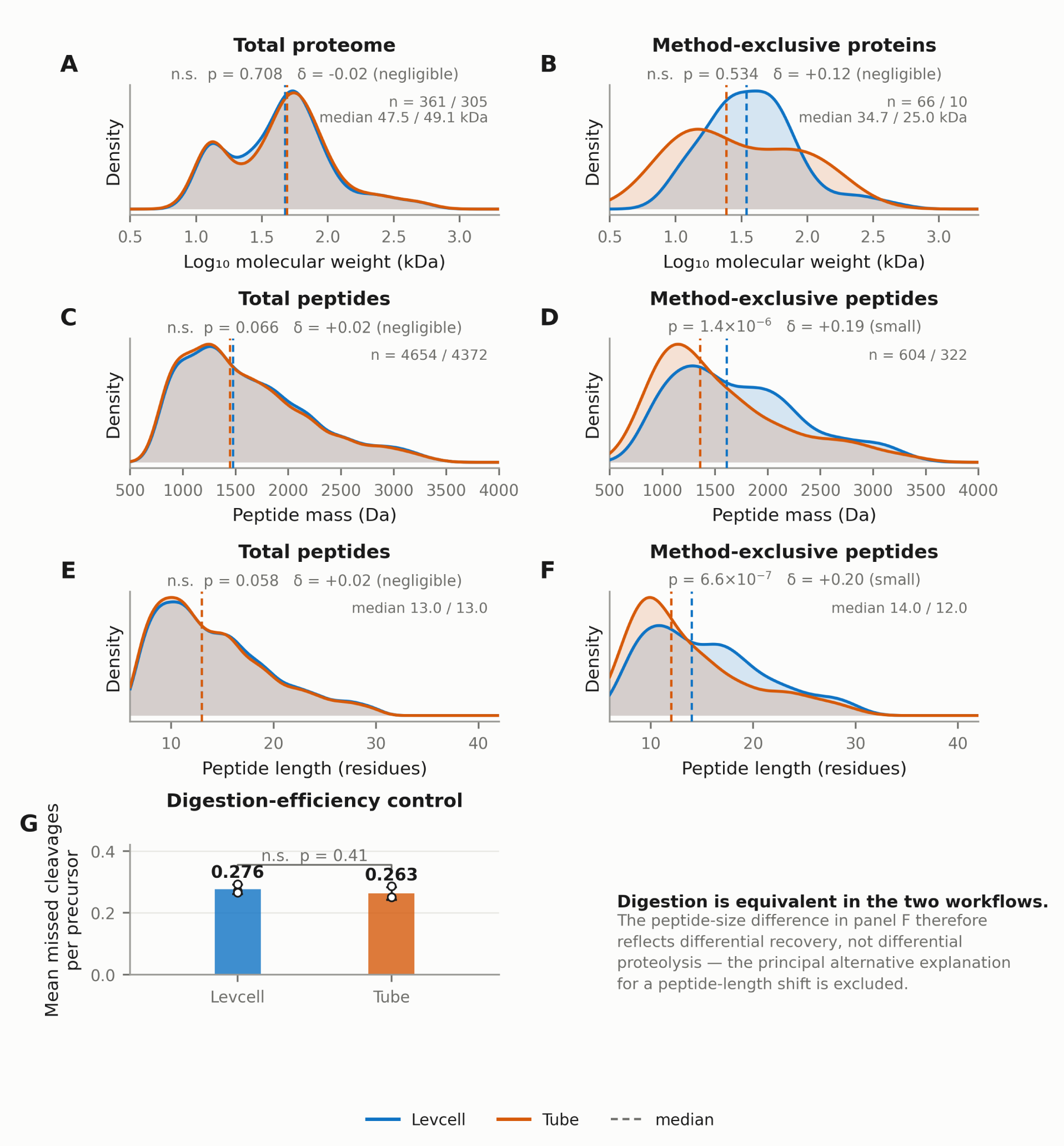
Size distributions of the recovered proteome and peptide pools. Kernel density estimates with dashed lines at the median; each panel reports a two-sided Mann–Whitney U test and Cliff’s delta as effect size. (A, B) Protein molecular weight for the (A) total proteome and (B) proteins exclusive to one workflow. (C, D) Peptide mass for the (C) total and (D) method-exclusive precursor pools. (E, F) Peptide length for the (E) total and (F) method-exclusive pools. (G) Mean missed tryptic cleavages per precursor per run, as a control for digestion efficiency; bars show the mean, error bars the standard deviation and open circles the individual runs. The equivalence in G establishes that the size difference in D and F reflects differential recovery rather than differential proteolysis.

Two controls bound the interpretation. First, the total precursor pools were indistinguishable between methods (median 13.0 residues in both; δ = +0.023, p = 0.058), so the size effect is confined to the differentially recovered fraction and does not shift the bulk digest. Second, and more importantly, the two workflows digested to the same extent: mean missed cleavages per precursor were 0.276 (levitation) versus 0.263 (tube) (Welch p = 0.41), and the proportion of precursors carrying at least one missed cleavage was 27.1% versus 25.7% (p = 0.38). Differential proteolysis — the most obvious alternative explanation for a peptide-length difference, given that a levitated droplet and a sealed tube present very different evaporative and interfacial environments — is therefore excluded.

At the protein level no size effect was resolvable. Median molecular weight was 47.5 kDa for the levitated proteome and 49.1 kDa for the tube proteome (p = 0.71), and the method-exclusive sub-proteomes did not differ significantly from each other (34.7 vs 25.0 kDa, p = 0.53), the tube-exclusive set comprising only ten proteins (Figure 2A, B). We therefore make no claim of a protein-level mass bias.

### No physicochemical selectivity is detectable

To test whether loss in tubes is mediated by hydrophobic or electrostatic interaction, we compared the intrinsic physicochemical properties of the recovered proteomes (Figure 3). None differed appreciably. Grand average of hydropathy was −0.28 for both total proteomes (p = 0.69), with method-exclusive medians of −0.32 (levitation) and −0.28 (tube) (p = 0.56). Hydrophobic residue frequency was 37.4% versus 37.2% in the total pools (p = 0.68). Isoelectric point medians were 6.07 in both total proteomes (p = 0.88), and 6.03 versus 5.67 in the exclusive subsets (p = 0.31).

**Figure 3.**
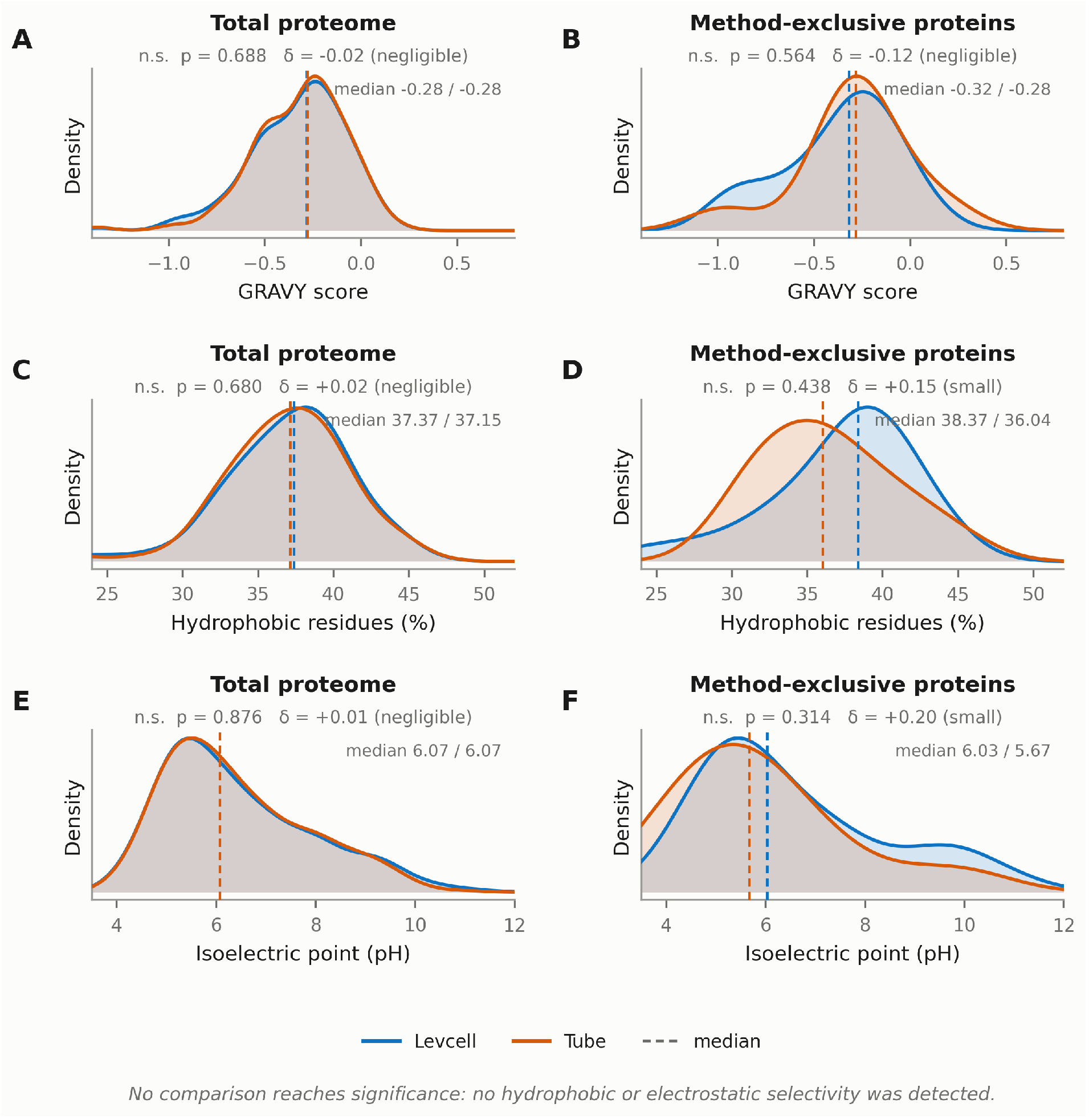
Physicochemical properties of the recovered proteomes, shown as kernel density estimates with medians marked. (A, B) Grand average of hydropathy, (C, D) hydrophobic residue frequency and (E, F) isoelectric point, for the total proteome (left) and the method-exclusive sub-proteomes (right). Properties were computed from UniProt canonical sequences. No comparison reaches significance; the study is underpowered to exclude moderate effects.

These null results are consistent with a sterically rather than chemically mediated loss process, but we note that with three replicates and a ten-protein tube-exclusive set the study is underpowered to exclude moderate hydrophobic or electrostatic contributions. We therefore state only that no such bias was detected, not that none exists.

### The rescued sub-proteome is enriched for canonical sEV luminal cargo

We next asked what kind of proteins container-less processing recovers. Over-representation analysis of the 66 levitation-exclusive protein groups was performed against the identified proteome as background — not against the whole genome, which inflates enrichment for any focused proteome — using KEGG, Reactome and MSigDB Hallmark gene sets, with Benjamini–Hochberg correction across all tested terms (Figure 5B).

The rescued set was strongly enriched for cytosolic macromolecular machinery: MYC targets (16 of 19 background genes, odds ratio 31.7, q = 5.6 × 10^-8^), RNA metabolism (18/25, OR = 15.7, q = 1.6 × 10^-7^), M phase and cell cycle (9/11 and 10/14, q *<* 0.001), driven by proteasome subunits (PSMA2, PSMA6, PSMB1, PSMB2), ribosomal proteins (RPL9, RPL17, RPS3A, RPS6), chaperonins (CCT2, CCT5) and het-erogeneous nuclear ribonucleoproteins (HNRNPA1, HNRNPA3, HNRNPK, HNRNPU) (Figure 5B). Note that the KEGG disease terms that also appear (for example Parkinson disease, Salmonella infection) share these same proteasome and ribosome members and are not independent findings. Cellular-component analysis gave a concordant picture, with enrichment for focal adhesion / cell-substrate junction (15/26, OR = 7.7, q = 6.9 × 10^-5^), intracellular non-membrane-bounded organelle (15/28, q = 1.7 × 10^-4^) and late endosome (4/5, q *<* 0.05) (Figure 4B). Proteasome, ribosome, chaperonin and hnRNP complexes are well documented sEV luminal cargo, and their recovery indicates that container-less processing preserves genuine low-abundance vesicular content rather than simply adding background. Results were essentially unchanged after removal of common contaminants.

**Figure 4.**
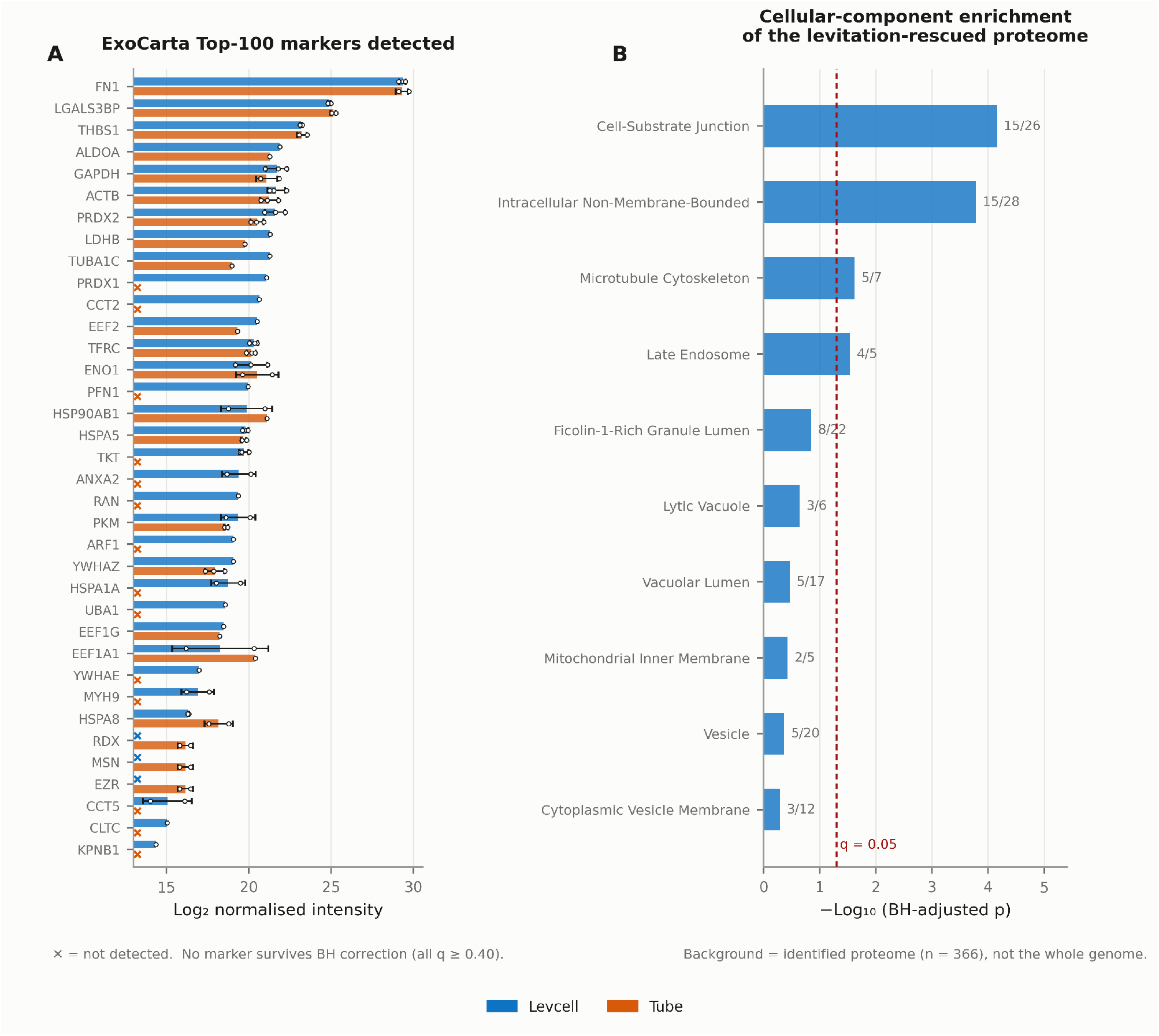
Recovery of canonical small-EV markers and cargo complexes. (A) Log_2_ normalised intensity of every ExoCarta Top-100 marker detected by either workflow, ordered by levitated abundance. Bars show the mean, error bars the standard deviation and open circles the individual replicates; × denotes a marker not detected in that workflow. Intensities are median normalised on the subset of proteins quantified in all six runs. No marker survives Benjamini–Hochberg correction, so no significance annotations are shown. (B) Gene Ontology cellular-component over-representation analysis of the 66 levitation-exclusive protein groups, using the identified proteome as background and one-sided Fisher’s exact tests with Benjamini–Hochberg correction across all tested terms. Labels give the overlap as k/K; the dashed line marks q = 0.05.

**Figure 5.**
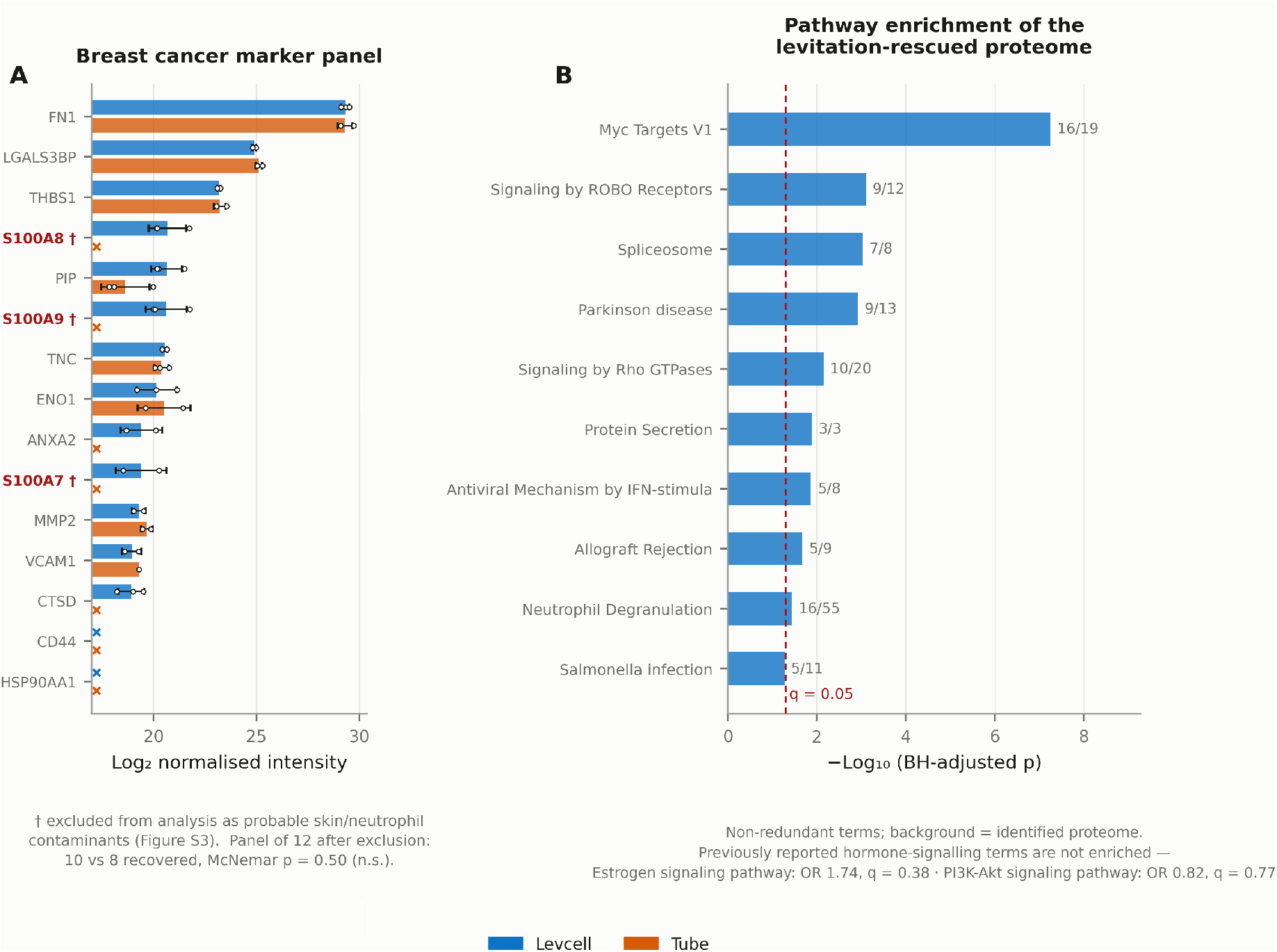
Breast cancer marker panel and functional enrichment. (A) Log_2_ normalised intensity of the 15-protein breast cancer panel; conventions as in Figure 4A. Members marked † and shown in red (S100A7, S100A8, S100A9) were excluded from analysis as probable skin- and neutrophil-derived contaminants (Supplementary Figure 3). CD44 and HSP90AA1 were not detected by either workflow. Of the remaining 12, ten were recovered under levitation and eight in tubes (exact McNemar p = 0.5). (B) Pathway over-representation analysis of the levitation-exclusive proteome against KEGG, Reactome and MSigDB Hallmark gene sets, computed as in Figure 4B. A non-redundant representative set is shown: terms sharing more than half their genes with a higher-ranked term are collapsed. KEGG disease terms share proteasome and ribosome members with the higher-ranked terms and are not independent findings.

Against the ExoCarta Top-100 reference set, levitated samples covered 33 markers and tubes 22 (Figure 1D, E; Figure 4A; Supplementary Figure 1). Fourteen markers — ANXA2, ARF1, CCT2, CCT5, CLTC, HSPA1A, KPNB1, MYH9, PFN1, PRDX1, RAN, TKT, UBA1 and YWHAE — were detected only under levitation. Three (EZR, MSN, RDX) were detected only in tubes; these constitute a single ezrin/moesin/radixin protein group, so the comparison at the level of marker-containing protein groups is 14 gained against one lost (exact McNemar p = 9.8 × 10^-4^). We note explicitly that this is a gain in absolute coverage rather than a proportional enrichment: markers constituted 9.1% of the levitated proteome and 7.2% of the tube proteome, a difference that is not significant (Fisher exact p = 0.40). Much of the marker gain therefore tracks the overall gain in depth.

### Marker-level abundance differences do not survive multiple-testing correction

We tested whether markers detected by both methods were also more abundant under levitation. This analysis is sensitive to normalisation, and we first note a methodological point of general relevance. Median normalisation applied across the full intensity column is biased when two conditions differ in detection depth: the deeper condition acquires a longer low-abundance tail, its column median falls, and median centring then shifts it upward. In our data this inflated the mean levitation-versus-tube difference among the 214 completely quantified proteins from +0.20 log_2_ (raw) to +0.72 log_2_. Normalising instead on the completely quantified subset removes the artifact, leaving a mean difference of +0.04 log_2_.

Using this unbiased normalisation, none of the 13 testable panel markers showed a significant abundance difference after Benjamini–Hochberg correction (smallest q = 0.396; PRDX2 log_2_FC +1.09, p = 0.073; PIP +2.01, p = 0.078). We therefore report no marker-level abundance advantage for container-less processing. The benefit we observe is one of detection — more proteins identified — not of quantitative gain for proteins already detected.

### Open-chamber processing carries a measurable contamination cost

Because the levitation chamber is open to its internal atmosphere and is served by a humidifier and robotic capillaries, we audited the recovered proteomes for the classic airborne contamination signature. Sixteen keratin-family protein groups were detected. Their summed share of total MaxLFQ signal was 3.10% in levitated samples (4.61%, 1.32%, 3.38%) versus 0.84% in tubes (1.39%, 0.66%, 0.48%) — a 3.7-fold higher burden, consistent in every replicate pair, although not significant with three replicates (p = 0.13). S100A8 and S100A9, which co-occur with keratins as a skin- and neutrophil-derived signature, were detected in all three levitated replicates and in none of the tube replicates; S100A7 was detected in two of three levitated replicates and no tube replicate (Supplementary Figure 3).

Two consequences follow. First, the coverage gain is not an artifact of this contamination: excluding keratins and S100A7/8/9 leaves 291.3 ± 20.6 versus 249.0 ± 6.6 protein groups (+17.0%, p = 0.059), and the paired asymmetry is unchanged. Second, proteins from this family cannot be interpreted as biological findings in this dataset. Accordingly we have excluded S100A7, S100A8, S100A9 and the keratins KRT5, KRT14 and KRT17 from all biomarker panels, notwithstanding their literature status as basal-like breast cancer markers.

### Breast cancer marker panel

Finally we assessed a curated panel of well-established breast cancer-associated proteins (Figure 5A). Two panel members, CD44 and HSP90AA1, were not detected by either workflow in this dataset and are therefore uninformative here. After removal of the contaminant-associated members, 10 of 12 panel proteins were recovered under levitation and 8 of 12 in tubes, with ANXA2 and cathepsin D (CTSD) detected only under levitation and none lost. This asymmetry does not reach significance (exact McNemar p = 0.5). Including the three S100 proteins would give 13 versus 8 (p = 0.063), still short of significance and, for the reasons above, not interpretable.

We therefore present this panel as an exploratory observation consistent with the general coverage gain, and make no claim that container-less processing preferentially rescues disease-specific signal. Establishing that would require biological replication across individual patients and a matched control group, neither of which the present pooled, single-sample design provides.

## Discussion

Adsorptive loss to container walls is a recognised constraint on low-input proteomics, and acoustic levitation offers a direct way to remove the interface at which that loss occurs. Applying an automated levitation platform to sEVs from breast cancer plasma, we find a reproducible increase in proteome coverage of roughly 18%, driven by a strongly asymmetric recovery of 66 additional proteins against 10 lost, with quantitative precision at least equal to that of tube-based processing.

The mechanistic evidence is consistent with size-dependent adsorptive loss, though it is more modest than we initially supposed. Peptides recovered only under levitation are longer and heavier than those recovered only in tubes, at a small effect size, and the total peptide pools are indistinguishable. Because a single protein yields many peptides, preferential loss of its longer sequences can reduce sequence coverage enough to drop the parent below the identification threshold, which would explain why a small peptide-level shift accompanies a substantial protein-level gain. The equivalence of missed-cleavage rates rules out the competing explanation that the levitated droplet simply digests differently. We could detect no hydropathy or charge selectivity, which is compatible with a steric mechanism but does not establish one.

Two observations argue against over-reading the result. First, once normalisation is performed in a way that is not biased by differential detection depth, no individual marker shows a significant abundance advantage. The benefit is in what is detected, not in how well it is quantified — a narrower but more defensible claim than the one we set out to test. Second, the open chamber has a cost: levitated samples carried roughly 3.7-fold more keratin signal, and the S100A7/A8/A9 group was exclusive to them. In a workflow whose purpose is to recover low-abundance analytes, an elevated airborne background is a material limitation, and it disproportionately affects exactly the low-abundance identifications the platform is meant to rescue. Practical mitigations — HEPA filtration of chamber make-up air, filtered humidifier water, and inclusion of a contaminant database in the search — are straightforward and should be standard for this platform.

The character of the rescued proteins is informative. Rather than a disease-specific signature, the levitation-exclusive set is dominated by ribosomal, proteasomal, chaperonin and RNA-binding complexes. These are established sEV luminal cargo, present at low abundance and therefore the first to be lost to a cumulative depletion mechanism. Their recovery is a coherent readout of reduced loss and, we would argue, better evidence of vesicular fidelity than the marker panels, which are small and depth-confounded.

### Limitations

This study has several constraints that bound its conclusions. The comparison rests on a single pooled plasma preparation processed in three parallel technical replicates; there is no biological replication across individual donors and no non-cancer control group, so no statement about disease specificity, and no biomarker claim, can be supported. Donor characteristics, including receptor status, are not available for a pooled sample, which is one reason we do not interpret hormone-signalling terms. Blood was collected into Streck tubes, which contain a formaldehyde-releasing preservative intended for cell-free DNA stabilisation and which may crosslink or modify proteins; the effect on EV proteome recovery is uncharacterised. EVs were isolated by polymer precipitation, which co-isolates lipoproteins and abundant plasma proteins, and our most intense protein is plasma fibronectin; the preparation should be regarded as sEV-enriched rather than pure. We did not perform orthogonal EV characterisation, so the study does not meet MISEV2023 reporting requirements in full. Samples were not reduced or alkylated, so cysteine-containing and disulfide-linked peptides are under-represented in both arms. Container contact is removed during digestion only; loading onto Evotips reintroduces a plastic interface, so the workflow mitigates rather than eliminates adsorptive loss. Finally, at three replicates per arm the study is underpowered for effects of the size we are attempting to resolve, and the primary count comparison is borderline (p = 0.053); the paired analyses should be regarded as the more reliable evidence.

## Conclusion

Automated container-less digestion in an acoustically levitated droplet increases the number of proteins identified from plasma sEV digests by approximately 18% relative to low-bind tubes, with equal or better reproducibility, and recovers a low-abundance sub-proteome of vesicular cargo complexes. The recovered peptides are modestly but reproducibly larger than those unique to tube processing, and this is not attributable to differential digestion, supporting size-dependent adsorptive loss as the underlying mechanism. Realising the approach in clinical proteomics will require contamination control appropriate to an open processing chamber, and validation in a design with biological replication.

## Materials and Methods

### Materials

Unless otherwise stated, all chemicals were purchased from commercial sources and used without further purification. HPLC-grade acetonitrile (ACN), formic acid (FA, ≥ 95%) and Tris base were from Sigma-Aldrich (MO, USA). Sequencing-grade modified trypsin (V5113) was from Promega (WI, USA). The Pierce BCA Protein Assay Kit was from Thermo Fisher Scientific (MA, USA). Trifluoroacetic acid (TFA) was from Chem-Impex International (IL, USA). The ExoQuick ULTRA EV Isolation kit was from System Biosciences (CA, USA).

### Human samples and ethics

Human plasma from breast cancer donors was provided by Dr Kent Hoskins’s clinical team at the University of Illinois Chicago. Samples were collected under an approved Institutional Review Board protocol (UIC Protocol #014-1078) in accordance with the Declaration of Helsinki, with written informed consent from all participants. Blood was collected into Streck tubes, processed to plasma by centrifugation, and stored at −80 °C. Plasma from donors was pooled prior to EV isolation; all downstream comparisons use this single pooled preparation.

### EV isolation

sEVs were isolated with the ExoQuick ULTRA kit according to the manufacturer’s protocol. Pooled plasma was thawed on ice and centrifuged at 3,000 × g for 15 min at 4 °C to remove cells and debris. The supernatant was transferred and centrifuged at 10,000 × g for 10 min at 4 °C. To 100 µL plasma, 27 µL ExoQuick precipitation reagent was added; the mixture was incubated on ice for 30 min and centrifuged at 3,000 × g for 10 min at 4 °C. The pellet was resuspended in 200 µL Buffer B, and 200 µL Buffer A was added. The purification column was washed and equilibrated with Buffer B (1,000 × g, 30 s). The EV mixture was loaded, incubated for 5 min at room temperature with gentle shaking, and eluted by centrifugation at 1,000 × g for 30 s. Eluate was aliquoted and stored at −80 °C. One aliquot was lysed by ice-bath sonication (1 min) and protein concentration determined by BCA assay.

### Levcell acoustic levitation platform

The Levcell device^17^ levitates droplets in a fully enclosed chamber with temperature and humidity control, giving stable reaction conditions and limiting evaporation. A camera measures droplet volume and adjusts humidifier output to compensate for detected changes. Three automated robotic arms deliver reagents from a 96-well plate to the levitated droplet through 250 µm inner-diameter capillary syringe pumps, permitting contact-free addition.

### EV sample preparation

Digestion was performed in parallel triplicate by acoustic levitation and in low-bind microcentrifuge tubes (Eppendorf). Each replicate used an EV eluate aliquot corresponding to 1 µg total protein. The Levcell chamber was held at 45 °C and 100% relative humidity using a deionised-water humidifier to prevent droplet evaporation.

For the container-less workflow, 3 µL of EV sample was dispensed and acoustically suspended by an automated robotic arm fitted with a 250 µm capillary, followed by contact-free addition of 2 µL digestion buffer. The final 5 µL droplet contained 50 mM Tris, 1 mM CaCl_2_ and trypsin at a 1:4 enzyme-to-protein ratio (w/w). Tube controls combined identical volumes of sample (3 µL) and buffer (2 µL) in low-bind tubes. Samples were not reduced or alkylated. All samples were digested simultaneously at 45 °C for 15 min and then loaded onto Evotips according to the manufacturer’s protocol.

## LC-MS/MS analysis

Digested peptides were separated on a 150 µm × 15 cm reversed-phase column packed with 1.9 µm C18 beads using an Evosep One LC system coupled to a Q Exactive HF mass spectrometer (Thermo Fisher Scientific). Samples were acquired in data-independent acquisition mode using the 44 min gradient, 30 samples-per-day method.

## Data processing

Raw files were converted to mzML (peak picking) with MSConvert and searched with DIA-NN v2.3.2 against a predicted spectral library generated from a UniProt reviewed human sequence database. Search parameters were: protease trypsin, cutting at K and R without proline restriction; up to 2 missed cleavages; peptide length 7–30 residues; precursor m/z 300–1800; precursor charge 1–4; fragment m/z 200–1800; Nterminal methionine excision enabled; no fixed modifications; variable modifications methionine oxidation (UniMod:35) and protein N-terminal acetylation (UniMod:1), maximum 2 per precursor; peptidoform scoring and RT profiling enabled. Match-between-runs was not used. Precursor and protein group identifications were filtered at 1% FDR; 381 protein groups passed a global q-value threshold of 0.01 across the six runs.

Marker-panel membership was evaluated across every gene listed in a protein group rather than the leading gene alone, because several reference markers are reported only within multi-gene groups (for example ACTB;ACTG1 and EZR;MSN;RDX). Where a set of markers falls in a single protein group, this is stated so that marker counts are not mistaken for independent identifications.

Downstream filtering, statistics and visualisation used a custom Python pipeline (Python 3.11; pandas, numpy, scipy, matplotlib). Protein intensities were log_2_ transformed and median normalised on the subset of proteins quantified in all six runs, rather than on the full intensity column, to avoid the depth-dependent bias described in the Results. A protein was required to have valid values in at least two of three replicates within a group to be evaluated in that group.

## Statistical analysis

Identification yields and per-run summary metrics were compared by two-sided Welch’s t-test. Because both workflows were applied to the same input material, presence/absence differences were assessed by exact McNemar test on discordant protein counts, and proportional enrichment of marker sets was assessed separately by Fisher’s exact test; we report both because they answer different questions. Distributions of peptide length, peptide mass, molecular weight, GRAVY, hydrophobic residue frequency and isoelectric point were compared by two-sided Mann–Whitney U test, with Cliff’s delta reported as effect size and interpreted as negligible (*<* 0.147), small (*<* 0.33), medium (*<* 0.474) or large. Per-marker abundance comparisons used Welch’s t-test with Benjamini–Hochberg correction across all markers tested; corrected q-values are reported and significance is claimed only at q *<* 0.05. Protein physicochemical properties were computed from UniProt canonical sequences.

## Functional enrichment analysis

Over-representation analysis of the levitation-exclusive protein set was performed with one-sided Fisher’s exact tests against KEGG (2021 Human), Reactome and MSigDB Hallmark gene sets, restricted to terms with at least two overlapping genes. The background was the set of genes identified in this experiment, not the whole genome. Benjamini–Hochberg correction was applied across all tested terms. No term filtering by disease keyword was applied. Because Reactome terms are strongly nested, a non-redundant representative set is shown; complete results are provided in Supplementary Table.

Subcellular localisation enrichment was assessed with the STRING database^22^.

## Supplementary Figures

**Supplementary Figure 1.**
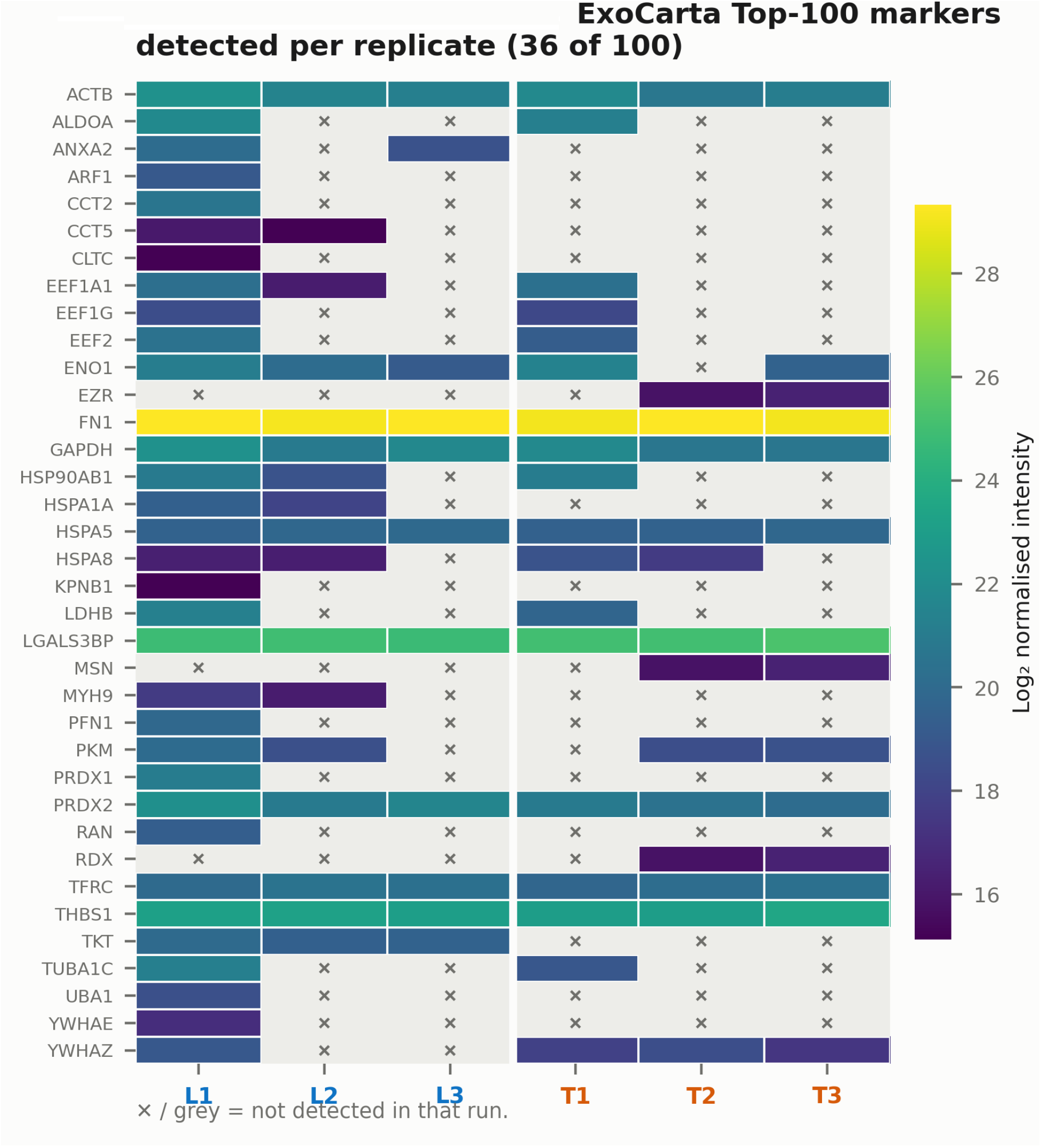
Per-replicate detection of ExoCarta Top-100 markers. Heat map of log_2_ normalised intensity for every reference marker detected in at least one run; grey cells marked *×* indicate no detection in that run.

**Supplementary Figure 2.**
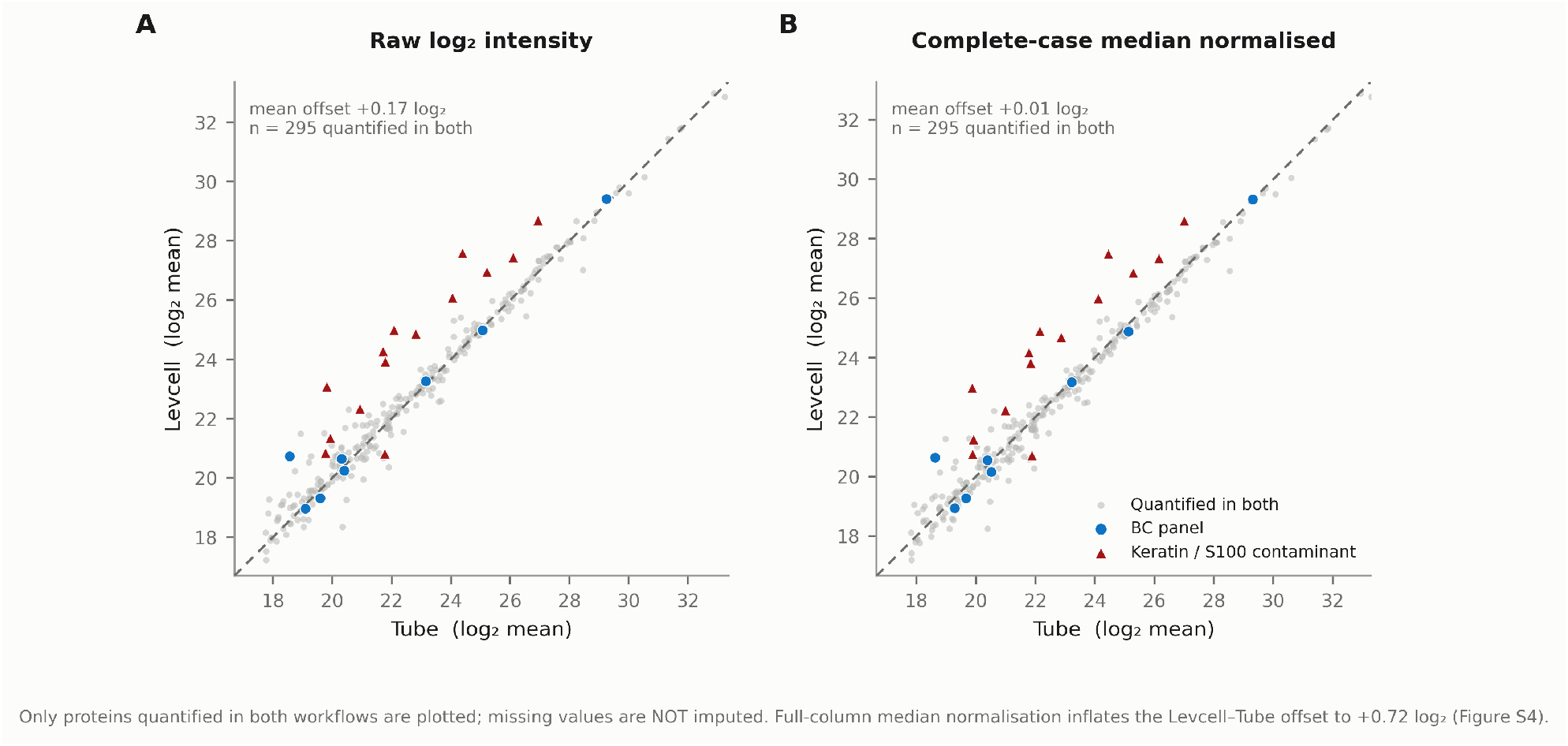
Protein-level abundance comparison, showing the effect of normalisation. Mean log_2_ intensity in levitated versus tube samples for the 295 proteins quantified in both workflows, (A) before and (B) after median normalisation on the completely quantified subset. Missing values are not imputed and undetected proteins are not plotted. The dashed line is the identity. Keratin and S100 contaminants (red triangles) lie systematically above the identity line in both panels.

**Supplementary Figure 3.**
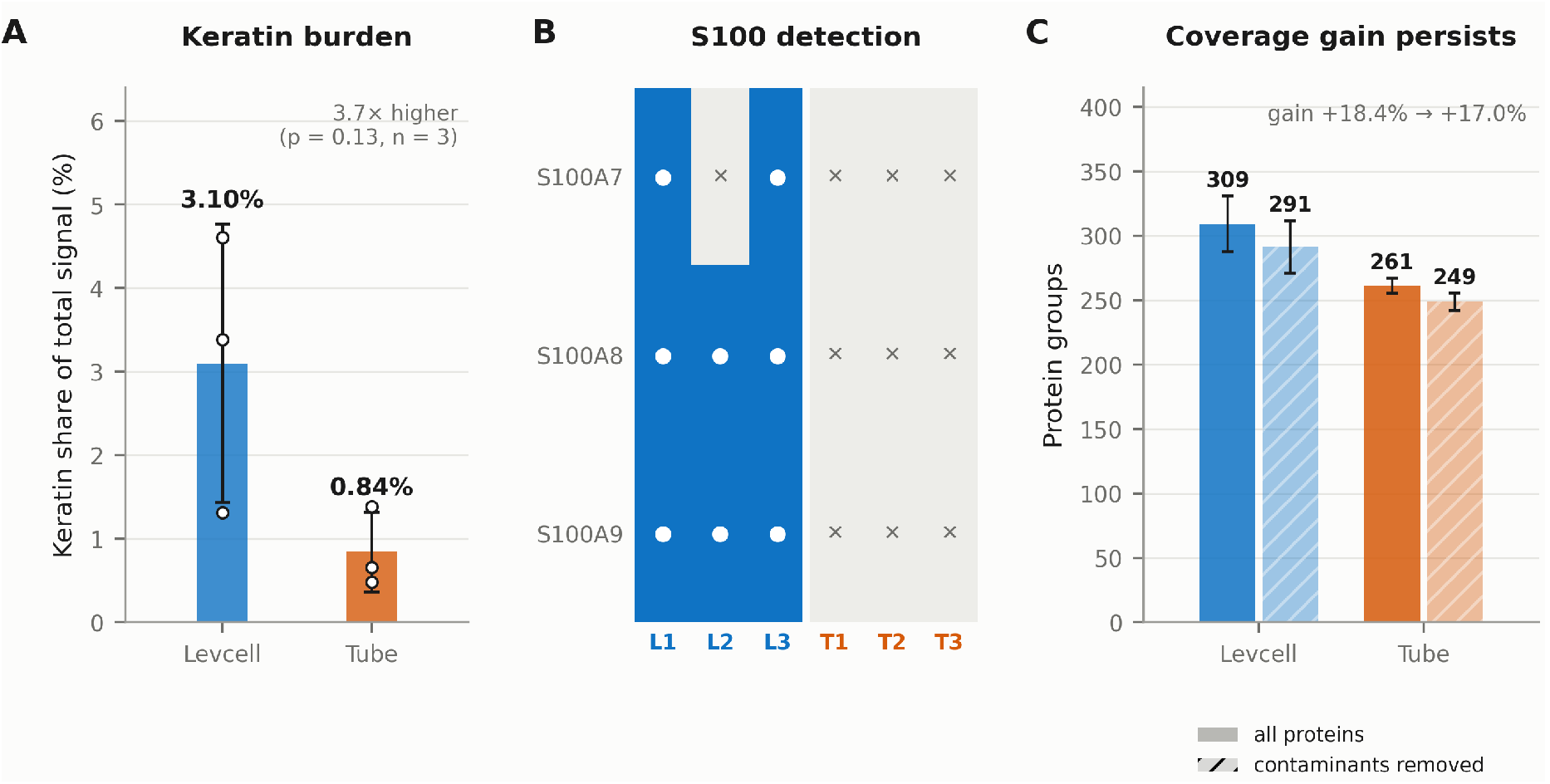
Airborne contamination audit. (A) Keratin-family share of the total summed MaxLFQ signal per run. (B) Per-run detection of S100A7, S100A8 and S100A9. (C) Mean protein groups identified before and after removal of keratins and S100A7/8/9, showing that the coverage gain is not an artifact of contamination.

**Supplementary Figure 4.**
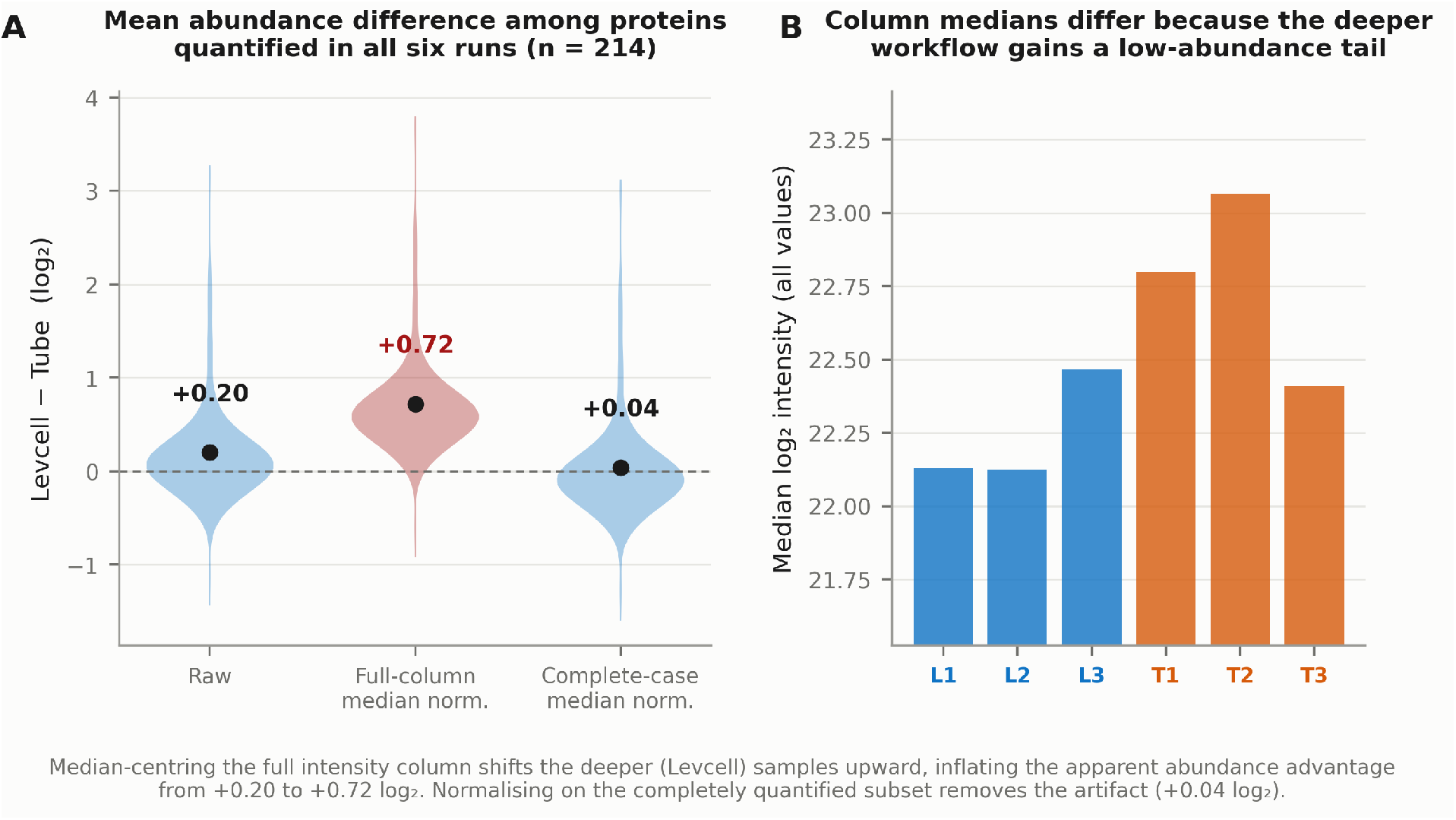
Normalisation diagnostic. (A) Distribution of the mean levitated-minus-tube log_2_ difference across the 214 proteins quantified in all six runs, under three normalisation schemes. Full-column median normalisation (red) inflates the difference from +0.20 to +0.72 log_2_; normalising on the completely quantified subset returns +0.04. (B) Median log_2_ intensity of each run computed over all quantified values, showing why full-column centring is biased when workflows differ in detection depth.

